# Insights into Origin and Evolution of α-proteobacterial Gene Transfer Agents

**DOI:** 10.1101/189738

**Authors:** Migun Shakya, Shannon M. Soucy, Olga Zhaxybayeva

**Affiliations:** Department of Biological Sciences, Dartmouth College, Hanover NH 03755, USA; Present address: Bioscience Division, Los Alamos National Laboratory, Los Alamos, NM 87544; Department of Computer Science, Dartmouth College, Hanover NH 03755, USA

**Keywords:** exaptation, domestication, horizontal gene transfer, bacterium-virus co-evolution, bacteriophage

## Abstract

Several bacterial and archaeal lineages produce nanostructures that morphologically resemble small tailed viruses, but, unlike most viruses, contain apparently random pieces of the host genome. Since these elements can deliver the packaged DNA to other cells, they were dubbed Gene Transfer Agents (GTAs). Because many genes involved in GTA production have viral homologs, it has been hypothesized that the GTA ancestor was a virus. Whether GTAs represent an atypical virus, a defective virus, or a virus co-opted by the prokaryotes for some function, remains to be elucidated. To evaluate these possibilities, we examined the distribution and evolutionary histories of genes that encode a GTA in the α-proteobacterium *Rhodobacter capsulatus* (RcGTA). We report that although homologs of many individual RcGTA genes are abundant across bacteria and their viruses, RcGTA-like genomes are mainly found in one subclade of α-proteobacteria. When compared to the viral homologs, genes of the RcGTA-like genomes evolve significantly slower, and do not have higher %A+T nucleotides than their host chromosomes. Moreover, they appear to reside in stable regions of the bacterial chromosomes that are generally conserved across taxonomic orders. These findings argue against RcGTA being an atypical or a defective virus. Our phylogenetic analyses suggest that RcGTA ancestor likely originated in the lineage that gave rise to contemporary α-proteobacterial orders *Rhizobiales, Rhodobacterales, Caulobacterales, Parvularculales, and Sphingomonadales,* and since that time the RcGTA-like element has co-evolved with its host chromosomes. Such evolutionary history is compatible with maintenance of these elements by bacteria due to some selective advantage. As for many other prokaryotic traits, horizontal gene transfer played a substantial role in the evolution of RcGTA-like elements, not only in shaping its genome components within the orders, but also in occasional dissemination of RcGTA-like regions across the orders and even to different bacterial phyla.

## 1 Introduction

Prokaryotes are hosts not only of their own genetic material, but also of mobile genetic elements, a broad class of entities that includes integrated viruses (prophages) (Frost et al. 2005). Traditionally, viruses are viewed as selfish genetic elements, but in some instances they can serve as vectors of horizontal gene transfer (HGT) (Touchon et al. 2017), an important driver of evolutionary success of many cellular lifeforms (Koonin 2016; Zhaxybayeva & Doolittle 2011). In other instances, prophage-like elements can provide immunity against other viruses (Canchaya et al. 2003), and therefore are beneficial to their host. While the evolutionary relationship between viruses and cellular lifeforms is still intensely debated, co-option of genes by both empires is likely frequent (Krupovic & Koonin 2017). For example, a few prokaryotic cellular functions that are associated with adaptations, such as cell-cell warfare, microbe-animal interactions, HGT, and response to environmental stress, are carried out using structures that resemble viral “heads” (e.g. McHugh et al. 2014; Sutter et al. 2008) or “tails” (e.g. Borgeaud et al. 2015; Shikuma et al. 2014). These structures appear to be widespread across archaea and bacteria (e.g. Böck et al. 2017; Sarris et al. 2014). In other instances, viral components appear to have originated from the cellular proteins (Krupovic & Koonin 2017). Such intertwined history of genes from cellular and non-cellular lifeforms resulted in a spectrum of genetic elements that, from the cellular host “point of view”, span from parasitic to benign to beneficial.

One class of such genetic elements is yet to be placed in this spectrum. Several unrelated bacterial and archaeal lineages are observed to produce particles that morphologically resemble viruses, yet appear to carry random pieces of host DNA instead of their own viral genome (Berglund et al. 2009; Bertani 1999; Humphrey et al. 1997; Marrs 1974; Rapp & Wall 1987). These particles were shown to transfer DNA between cells through a transduction-like mechanism (Lang et al. 2012; McDaniel et al. 2010), and that’s why they were dubbed “Gene Transfer Agents” (GTAs) (Marrs 1974). Since the transferred DNA may benefit the recipient cells, GTAs were postulated to be viruses that were "domesticated” by prokaryotes (Bobay et al. 2014) and maintained by them as a mechanism of HGT (Lang et al. 2012). Yet, the details of such co-option event as well as impact of the GTA-mediated HGT remain to be deciphered, and alternatives, such as GTA being instead a defective or atypical virus, need to be evaluated.

The laboratory studies of *Rhodobacter capsulatus*’ GTA (or RcGTA for short) show that this element neither has a typical genome of a bacterial virus nor acts like a typical lysogenic virus. Its genome is distributed among five loci dispersed across the *R. capsulatus* chromosome (Hynes et al. 2016 and **Figure 1A**). Seventeen of the genes are found in a single locus that has a genomic architecture typical of a siphovirus (Lang & Beatty 2007). Most of the genes in the locus encode proteins involved in head and tail morphogenesis of the RcGTA particle and are thus referred to as the ‘head-tail’ cluster (after Lang et al. 2017). The remaining four loci contain seven genes shown to be important for RcGTA production, release, and DNA transfer (Fogg et al. 2012; Hynes et al. 2012, 2016; Westbye et al. 2015). Even if RcGTA would preferentially package the regions of DNA that correspond to its own genome, the small head size of RcGTA particle can accommodate only ~1/5 of the genome (Lang et al. 2017), and therefore RcGTA can propagate itself only with the division of the host cell. Expression of the RcGTA genes, as well as production and release of RcGTA particles can be triggered by phosphate concentration (Leung et al. 2010; Westbye et al. 2013), salinity (McDaniel et al. 2012), and quorum sensing (Brimacombe et al. 2013; Schaefer et al. 2002). The latter is regulated via CckA-ChpT-CtrA phosphorelay (Leung et al. 2012; Mercer & Lang 2014), a widely used bacterial signaling network that also controls cell cycle (Chen & Stephens 2007; Mann et al. 2016) and flagellar motility (Zan et al. 2013). These observations suggest that RcGTA is under control of the bacterial host and is well-integrated with the host’s cellular systems. When did such integration occur? Could RcGTA still represent a selfish genetic element? Through phylogenomic analyses of a much more extensive genomic data set, we show that this α-proteobacterial GTA is a virus-related element whose genome evolves differently from what is expected of a typical bacterial virus. We also infer that RcGTA-like element likely originated in an ancestor of an α-proteobacterial subclade that gave rise to at least five contemporary taxonomic orders. Since that time, the genomes of these elements were shaped by both co-evolution with the bacterial hosts and HGT.

**Figure 1:**
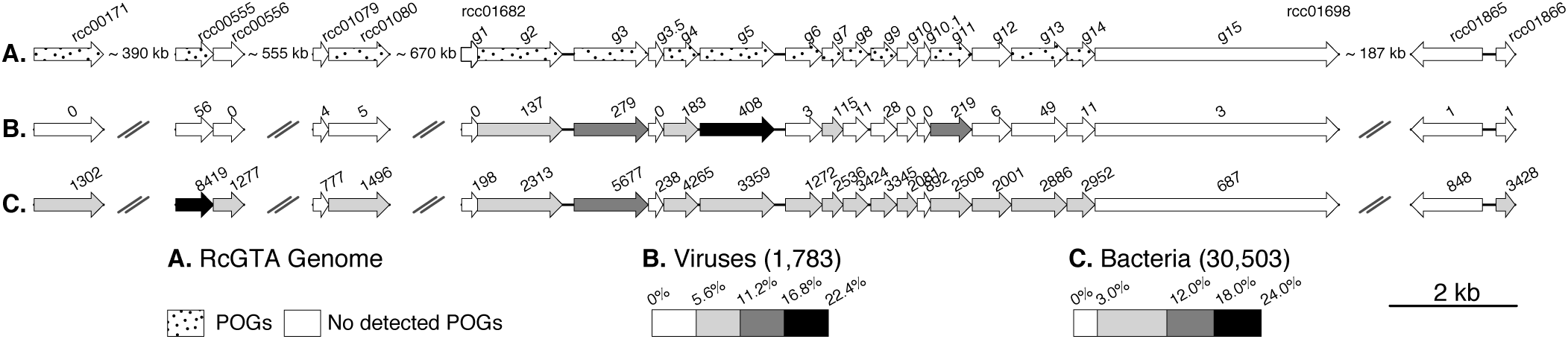
**RcGTA genome and distribution of its homologs in viral and bacterial genomes. Panel A. The RcGTA genome architecture.** The RcGTA genome consists of 24 genes scattered across five loci in the *R. capsulatus* SB1003 genome. The genes are represented by arrows. The majority of the ‘head-tail’ cluster genes (*R. capsulatus* SB1003 locus tags *rcc01682*-*rcc01698*; also known as *g1*-*g15*) encode genes involved in head and tail morphogenesis, *rcc00171* (*tsp*) - a tail spike protein (Hynes et al. 2016), *rcc00555* and *rcc00556* - endolysin and holin (Fogg et al. 2012; Hynes et al. 2012; Westbye et al. 2013), *rcc01079* (*ghsA*) and *rcc01080* (*ghsB*) - a head spike (Westbye et al. 2015), and *rcc01865* and *rcc01866* - putative regulatory elements (Hynes et al. 2016). The GenBank functional annotations of the genes are listed in **Supplementary Table S4**. Genes with similarity to gene families in the POG database (Kristensen et al. 2013) are shaded and their assigned POG categories are listed in **Supplementary Table S4. Panel B. Presence of RcGTA gene homologs in 1,783 viral genomes. Panel C. Presence of RcGTA gene homologs in 30,503 bacterial genomes.** The numbers above arrows represent the total number of detected homologs for each RcGTA gene, while the percent of viral (**panel B**) bacterial (**panel C**) genomes in which these homologs are found is depicted in shades of gray (see figure inset for scale). Genomic regions were visualized using R package genoplotR (Guy et al. 2010).

## 2 Methods

### Detection of RcGTA homologs in viruses and bacteria

Twenty-four genes from the RcGTA genome and their homologs from *Rhodobacterales* were used as queries in BLASTP (E-value < 0.001; query and subject overlap by at least 60% of their length) and PSI-BLAST searches (E-value < 0.001; query and subject overlap by at least 40% of their length; maximum of 6 iterations) against viral and bacterial databases using BLAST v. 2.2.30+ (Altschul et al. 1997). The viral database consisted of 1,783 genomes of dsDNA bacterial viruses extracted from the RefSeq database (release 76; last accessed on July 1, 2016) using “taxid=35237 AND host=bacteria” as a query. Bacterial RefSeq database (release 76; last accessed on June 29, 2016) and 255 completely sequenced α-proteobacterial genomes (**Supplementary Table S1**; genomes downloaded from ftp://ftp.ncbi.nlm.nih.gov/genomes/Bacteria/ on November 1, 2015) served as two bacterial databases. The detected RefSeq homologs were mapped to their source genomes using "release76.AutonomousProtein2Genomic.gz" file (ftp.ncbi.nlm.nih.gov/refseq/release/release-catalog/, last accessed on June 29, 2016).

### Assignment of genes to viral gene families

All annotated protein-coding genes from 255 α-proteobacterial genomes were used as queries in BLASTP searches (E-value < 0.001; query and subject overlap by at least 40% of their length) against the POG (Phage Orhologous Group) database (Kristensen et al. 2013), which was downloaded from ftp://ftp.ncbi.nlm.nih.gov/pub/kristensen/thousandgenomespogs/blastdb/ in February 2016. The genes were assigned to a POG family based on the POG affiliation of the top-scoring BLASTP match.

### Identification of RcGTA homologs adjacency in the bacterial genomes

Relative positions of RcGTA homologs in each genome were calculated from genome annotations in the GenBank feature tables downloaded from ftp.ncbi.nlm.nih.gov/genomes/all/ on June 29, 2016. Initially, a region of the genome was classified as putative RcGTA-like region if it had at least one RcGTA homolog. The identified regions were merged if two adjacent regions were separated by < 15 open reading frames (ORFs). If the resulting merged region contained at least nine RcGTA homologs, it was designated as a large cluster (LC). The remaining regions were labeled as small clusters (SC).

### Examination of viral characteristics of RcGTA-like genes and regions

Prophage-like regions in 255 α-proteobacterial genomes were detected using PhiSpy v. 2.3 (Akhter et al. 2012) with “-n=5” (at least 5 consecutive prophage genes), “-w=30” (a window size of 30 ORFs), and “-t=0” (“Generic Test Set”) settings. An RcGTA-like region was classified as a putative prophage if it overlapped with a PhiSpy-detected prophage region.

Presence of viral integrases was assessed within all prophage-like and RcGTA-like regions, as well as among 5 ORFs upstream and downstream of the regions. The integrase homologs were identified using the hmmsearch program of the HMMER package v. 3.1b2 (Eddy 1998; E-value < 10^-5^), with HMM profiles of integrase genes from the pVOG database 2016 update (Grazziotin et al. 2017) as queries (VOG# 221, 275, 375, 944, 2142, 2405, 2773, 3344, 3995, 4650, 5717, 6225, 6237, 6282, 6466, 7017, 7518, 8218, 8244, and 10948) and 255 α-proteobacterial genomes as the database.

For both large and small clusters, %G+C of the corresponding nucleotide sequence (defined as DNA sequence between the first and last nucleotide of the first and last RcGTA homolog of the region, respectively) was calculated using in-house Python scripts. The relative difference in GC content of the RcGTA-like region and the whole host genome was calculated by taking the difference in their %G+C and dividing it by the %G+C of the host genome. This correction allowed us to compare changes in the GC content of the RcGTA-like regions among genomes with variable GC contents. To account for possible heterogeneity of GC content within each genome, 100 genomic regions of the same size were randomly selected from the genome, their relative GC content calculated as described above and compared to that of the RcGTA-like region(s) using pairwise t-test.

Pairwise phylogenetic distances (PPD) among RcGTA homologs in viruses, and large and small clusters were calculated in RAxML v. 8.1.3 (Stamatakis 2014) using ‘-f x’ and ‘-m PROTGAMMAAUTO’ options. The latter choice selects the best amino acid substitution model and uses Γ distribution with four discrete rate categories to correct for among site rate variation (Yang 1994).

### Identification of gene families

Sequences of protein-coding genes in 255 α-proteobacterial genomes were subjected to all-against-all BLASTP searches (v. 2.2.30+, E-value < 0.0001), in which reciprocal top-scoring BLASTP matches were retained. The bit scores of the matches were converted to pairwise distances, which were used to form gene families via Markov clustering with the inflation parameter set to 1.2 (van Dongen 2000), as implemented in OrthoMCL v. 2.0.9 (Li 2003). The gene presence/absence in the resulting 33,048 gene families was used to as a proxy of gene family conservation within and across α-proteobacteria.

### Characterization of immediate gene neighborhoods of small and large clusters

ORFs without detectable similarity to RcGTA genes, but found immediately upstream, downstream, or within ‘head-tail’ cluster, were classified based on (a) conservation within α-proteobacterial genomes and (b) potential viral origin. The proportion of α-proteobacterial genomes that have an ORF homolog was used as a proxy for conservation. The ORF was designated as viral if it was assigned to a POG.

### Reference phylogeny of α-proteobacteria

From the list of 104 genes families used to reconstruct α-proteobacterial phylogeny (Williams et al. 2007), 99 gene families that are present in 80% of the 255 α-proteobacterial genomes were selected (**Supplementary Table S2**). For each gene family, homologs from *Geobacter sulfurreducens* PCA, *Ralstonia solanacearum* GMI1000, *Chromobacterium violaceum* ATCC 12472, *Xanthomonas axonopodis* pv. citri str. 306*, Pseudomonas aeruginosa*PAO1*, Virbrio vulnificus* YJ016*, Pasteurella multocida* subsp *multocida* str Pm70, *Escherichia coli* str. K12 substr. DH10B were added as an outgroup. The amino acid sequences of the resulting gene set were aligned using MUSCLE v. 3.8.31 (Edgar 2004). The best substitution model (listed in **Supplementary Table S2**) was selected using *ProteinModelSelection.pl* script downloaded from http://sco.h-its.org/exelixis/resource/download/software/ProteinModelSelection.pl in August 2016. Among site rate variation was modeled using Γ distribution with four rate categories (Yang 1994). All gene sets were combined into a single dataset with 99 data partitions (one per gene.) The maximum likelihood phylogenetic tree was reconstructed in RAxML v. 8.1.3 (Stamatakis 2014), using the best substitution model for each partition and 20 independent tree space searches.

One hundred bootstrap samples were analyzed as described above, and the bootstrap support values > 60% were mapped to the maximum likelihood tree. Reference phylogeny of only LC-containing taxa was obtained by pruning taxa that only have SCs.

### Phylogenetic analysis of the large-cluster gene families

Since phylogenetic histories of many RcGTA homologs are poorly resolved in *Rhodobacterales* (Hynes et al. 2016), amino acid sequences of LC gene families were combined into one “LC-locus” dataset in order to increase the number of phylogenetically informative sites. To ensure that such concatenation does not result in a mixture of genes with different evolutionary histories, phylogenetic trees reconstructed from the alignments of individual LC genes and of the concatenated LC-locus were compared. Amino acid sequences of gene families consisting only of LC homologs from α-proteobacterial genomes were aligned using MUSCLE v. 3.8.31 (Edgar 2004). The individual gene family alignments were concatenated into the “LC-locus” alignment. Since many LCs do not have homologs of all

RcGTA ‘head-tail’ genes, in the concatenated alignment the absences were designated as missing data (Stamatakis 2014). Additionally, “taxa-matched” LC-locus alignments (*i.e.* concatenated alignments pruned to contain taxa found only in a specific LC gene family) were created. Maximum likelihood trees were reconstructed from the concatenated and individual LC gene alignments, as well as from the taxa-matched LC-locus alignments, using RAxML v. 8.2.9 (Stamatakis 2014) under the LG+Γ substitution model (Le & Gascuel 2008; Yang 1994). The LG model was determined as the best substitution model for each individual gene set using the PROTGAMMAAUTO option of RAxML v. 8.2.9 (Stamatakis 2014). For each LC gene, consensus trees were reconstructed from 10 bootstrap samples of taxa-matched LC-locus alignments. These consensus trees were compared to 100 bootstrap sample trees for the corresponding LC gene. The congruence between the phylogenies was measured using relative “<u>T</u>ree <u>C</u>ertainty <u>A</u>ll” (TCA) values (Salichos et al. 2014), as implemented in RAxML v. 8.2.9 (Stamatakis 2014). The TCA values were classified into “strongly conflicting” (−1.0 ≤ TCA ≤ −0.7), “moderately conflicting” (-0.7 < TCA < 0.7) and “not conflicting” (0.7 ≤ TCA ≤ 1.0) categories. None of the individual LC gene phylogenies had strongly conflicting TCA values, although 6 out of 17 LC gene phylogenies had moderately conflicting values (**Supplementary Table S3**). Since an uncertain position of just one taxon can affect support values of multiple bipartitions (Aberer et al. 2013) and TCA values calculated from them, a Site Specific Placement Bias (SSPB) analysis (Berger et al. 2011) was carried out, as implemented in RAxML v. 8.2.9 (Stamatakis 2014). The LC-locus phylogeny and the LC-locus alignment were used as the reference tree and the input sequence alignment, respectively. The sliding window size was set to 100 amino acids. The SSPB analysis revealed that for 15 of the 17 genes only three nodes, on average, separate the optimal positions of each taxon in the gene and LC-locus phylogenies, indicating largely compatible evolutionary histories of individual LC genes and of their combination (**Supplementary Figure S1**). Consequently, we decided to use the combined, taxa-matched LC-locus alignment for further phylogenetic analyses.

To assess the similarity of the evolutionary histories of LC loci and of their hosts, the topologies of the reference α-proteobacterial tree were compared to 100 bootstrap sample trees reconstructed from the taxa-matched LC-locus alignment assembled as described above. The congruence was quantified using Internode <u>C</u>ertainty (IC) values (Salichos et al. 2014), as implemented in RAxML v. 8.2.9 (Stamatakis 2014). The IC values were classified into “strongly conflicting” (-1.0 ≤ IC ≤ −0.7), “moderately conflicting” (-0.7 < IC < 0.7) and “not conflicting” (0.7 ≤ IC ≤ 1.0) categories.

### Conservation of gene neighborhoods across phylogenetic distance

Forty ORFs upstream and downstream of the ATP synthase operon, the ribosomal protein operon, and the LCs were extracted from 87 α-proteobacterial genomes with at least one LC. The same procedure was performed for a transposase from IS3/IS911 family, which was detected in 18 of the 87 genomes. Each ORF was assigned to a gene family, and the pairwise conservation of the corresponding flanking regions was calculated as a proportion of gene families shared between a pair of genomes. PPDs of 16S rRNA genes were used as a proxy for time *t*. 16S rRNA genes were aligned against the GreenGenes database (Desantis et al. 2006; last accessed in August 2016) using mothur v. 1.35.1 (Schloss et al. 2009). The PPDs were calculated from the alignment using RAxML v. 8.1.3 (Stamatakis 2014) under GTR+Γ substitution model. Based on *model 3* from Rocha (2006), the gene neighborhood decay was modeled as *p^t^,* where *p* is the probability of the 40 ORFs to remain in the same genomic region and *t* is time. This model assumes that every gene has equal probability of separating from the region. The parameter *p* was estimated, and the fit between the model and data was assessed using non-linear least squares method (Bates & Watts 1988), as implemented in the *nls* R function.

### Examination of genes that flank RcGTA-like clusters in non-α-proteobacterial genomes

Amino acid sequences of five ORFs upstream and downstream of the RcGTA-like clusters in the genomes of actinobacteria *Streptomyces purpurogeneiscleroticus* NRRL B-2952 (GenBank accession number LGEI00000000.1;Ju et al. 2015) and *Asanoa ferruginea* NRRL B-16430 (LGEJ00000000.1; Ju et al. 2015), a γ-proteobacterium *Pseudomonas bauzanensis* W13Z2 (JFHS00000000.1; Wang et al. 2014), and a cyanobacterium *Scytonema millei* VB511283 (JTJC00000000.1; Sen et al. 2015) were searched against the database of protein-coding genes of 255 α-proteobacterial genomes using BLASTP (E-value < 0.001; query and subject overlap by at least 60% of their length). Three genes that flank LCs in *P. bauzanensis*, *S. millei* and a few α-proteobacteria (*kpdE*, *pbpC*, *dnaJ*) were used as queries to detect their homologs in other cyanobacteria and γ-proteobacteria in BLASTP searches of γ-proteobacteria and cyanobacteria in the *nr* database (<u>https://blast.ncbi.nlm.nih.gov/Blast.cgi</u>, last accessed on October 31, 2016; E-value < 0.001; query and subject overlap by at least 60% of their length). For each of the three gene sets, four homologs from each identified taxonomic group were selected and combined with all available homologs from LC-containing α-proteobacteria. The amino acid sequences of the combined data sets were aligned using MUSCLE v. 3.8.31 (Edgar 2004). Maximum likelihood trees were reconstructed using FastTree v. 2.1.8 (Price et al. 2010) under JTT+Γ substitution model. The trees were visualized and annotated using EvolView <u>(He et al. 2016; http://evolgenius.info/evolview,</u> last accessed on November 13, 2016).

### Identification of CRISPR/*Cas* defense systems

Spacers of the Clustered Regularly Interspaced Short Palindromic Repeats (CRISPRs) were identified by scanning 255 α-proteobacterial genomes for both CRISPR leader sequences and repeat structures using CRISPRleader v. 1.0.2 (Alkhnbashi et al. 2016) and PILER-CR v. 1.06 (repeat length between 16 and 64; spacer length between 8 and 64; minimum array length of 3 repeats; minimum conservation of repeats 0.9; minimum repeat length ratio 0.9; minimum spacer length ratio 0.75) (Edgar 2007), respectively. CRISPR-associated proteins were identified using the hmmsearch program of the HMMER package v. 3.1b2 (Eddy 1998; E-value < 10^-5^), with TIGRFAM families of the CRISPR-associated proteins as queries (TIGRFAM # 287, 372, 1573, 1587, 1596, 1863, 1865, 1868, 1869, 1876, 1907, 2562, 2589, 2590, 2593, 2621, 3158, 3637, 3638, 3639, 3640, 3641, 3983; Selengut et al. 2007), and protein-coding genes of 255 α-proteobacterial genomes as database. Genomes were designated to have putative functional CRISPR/*Cas* systems if both CRISPR spacer arrays and at least three CRISPR-associated proteins from the same CRISPR class (Makarova et al. 2015) were present. Spacer sequences from these putatively functional CRISPR systems were used as BLASTN (v. 2.4.0+; E-value < 10; task = tblastn-short) queries against all gene sequences in LCs.

### Identification of decaying RcGTA ‘head-tail’ cluster homologs

The nucleotide sequences of LC and SC regions were extracted from each genome and translated in their entirety into all six reading frames. RcGTA ‘head-tail’ cluster genes were used as BLASTP (v. 2.4.0+, E-value ≤ 0.001) queries in searches against the database of translated LC and SC regions. Homologous sequences within LC and SC regions that did not match the annotated ORFs were designated as putatively decaying RcGTA genes.

## 3 Results

### Many RcGTA genes share evolutionary history with viruses

Consistent with the earlier proposed hypothesis that RcGTA is an element originated from a virus (Lang & Beatty 2000), 14 out of 24 RcGTA genes can be assigned to a POG (**Figure 1A** and **Supplementary Table S4**), and 18 out of the 24 genes have at least one readily detectable homolog in available genomes of *bona fide* bacterial viruses (**Figure 1B**). However, none of the bacterial viruses in RefSeq database contained a substantial number of the RcGTA homologs (at most eight of them in *Rhizobium* phage 16-3 [GenBank Acc. No. DQ500118.1]), and no α-proteobacterial genomes with CRISPR systems contained significant matches between CRISPR spacers and RcGTA genome, suggesting that the presumed progenitor virus is either extinct or remains unsampled. Collectively, RcGTA-like genes are unevenly represented across viral genomes (**Figure 1B**). Among the five most abundant are homologs of *rcc001683* (*g2*), *rcc001684* (*g3*), *rcc001686* (*g4*) and *rcc001687* (*g5*), which encode terminase, portal protein, prohead protease, and major capsid protein, respectively.

Their widespread occurrence is not surprising. First, these genes correspond to the so-called “viral hallmark genes”, defined as genome replication and virion formation genes with a wide distribution among diverse viral genomes (Iranzo et al. 2016; Koonin et al. 2006). Second, *g3*, *g4* and *g5* belong to HK97 family, and HK97-like genes are common in Caudovirales (Iranzo et al. 2017) - one of the most abundant viral groups (Cobián Güemes et al. 2016). However, 13 of the RcGTA genes are either sparsely represented (found in 10 or fewer viral genomes; 7 genes) or not detected at all (6 genes) in viruses. These genes may be fast-evolving viral genes, auxiliary (non-hallmark) viral genes, which could be specific to the viral lineage that gave rise to RcGTA, non-viral (cellular) genes, or simply be a result of undetected homology due to the paucity of viral genomes in GenBank. The latter possibility cannot be ignored, especially given that (i) three of the 13 genes are assigned to a POG (**Figure 1A**), and therefore are likely of viral origin, and (ii) only 94 of the 1,783 screened bacterial viruses (5%) have α-proteobacteria as their known host. Since about half of bacterial genomes are estimated to contain at least one prophage in their genome (Touchon et al. 2016), inclusion of RcGTA homologs from bacterial genomes could reduce the possibility of undetected homology due to lack of available viral genomes. Therefore, we looked for RcGTA homologs in all bacterial RefSeq records, and in 255 completely sequenced α-proteobacterial genomes in particular (**Supplementary Table S1**).

### Homologs of RcGTA genes are abundant in bacteria, but RcGTA-like genomes are mainly found in class α-proteobacteria

Every RcGTA gene has at least 198 homologs in RefSeq database (**Figure 1C**), and therefore even the 13 “rare” genes are relatively abundant in bacterial genomes. Moreover, six of the 13 genes (*rcc00171*, *rcc00556*, *rcc01688* [*g6*], *rcc01692* [*g10*], *rcc001695* [*g12*], and *rcc01866*), including three assigned to POGs, are found in more than 1000 bacterial genomes (**Figure 1C**). Therefore, it is unlikely that the 13 rare genes are auxiliary viral genes specific to the viral lineage that gave rise to RcGTA. However, a larger number of genomes of α-proteobacterial viruses is needed to identify if these genes are of viral or cellular origin.

Collectively, homologs of RcGTA genes are found within 13 bacterial phyla (**Figure 1C** and **Supplementary Figure S2a**). Interestingly, 84% and 35% of the identified homologs belong to the phylum Proteobacteria and to its class α-proteobacteria, respectively.

Furthermore, many α-proteobacterial genomes contain more ‘head-tail’ RcGTA homologs than any available viral genome, and these homologs are often clustered (**Supplementary Figure S2b**). In contrast, most other taxa have only few RcGTA homologs that are scattered across the genome (**Supplementary Figures S2b** and **S2c**). Since in the RcGTA genome the ‘head-tail’ genes are in one locus (**Figure 1A**) and are presumed to be co-transcribed (Kuchinski et al. 2016), functional GTAs of this family might be restricted mainly to α-proteobacteria.

Within α-proteobacteria, homologs of RcGTA genes are detectable in 197 out of 255 examined genomes (77%). These homologs are distributed across eight α-proteobacterial orders with at least one available genome and two unclassified α-proteobacterial genera (**Figure 2**). While 13 out of 17 ‘head-tail’ genes are abundant across α-proteobacteria (and across Bacteria in general), only two of the genes from the remaining four loci (*rcc00555* and *rcc01866*) are frequently detected in bacterial genomes (**Figure 1C** and **Figure 2**). Since within *Rhodobacterales* genes outside of ‘head-tail’ cluster evolve faster than their ‘head-tail’ counterparts (Hynes et al. 2016), we conjecture that many α-proteobacterial homologs of the former genes were not detected in our searches. Therefore, for the subsequent investigations we focused mostly on the analyses of the ‘head-tail’ locus genes.

**Figure 2:**
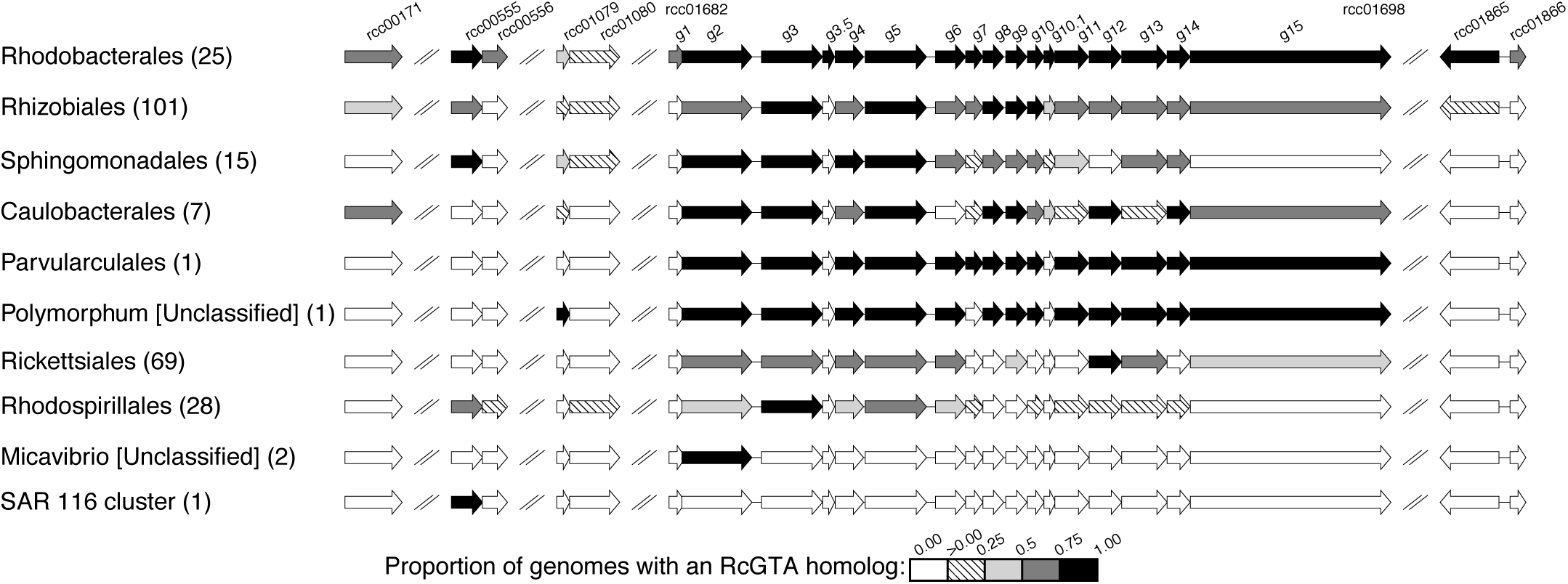
**Presence of RcGTA-like genes in α-proteobacterial genomes.** Each line represents the distribution of genes in the RcGTA genome within a taxonomic group. The number of surveyed genomes in a taxonomic group is listed in parentheses. The shades of gray represent the proportion of these genomes in which the RcGTA homologs are found (see figure inset). Genomic regions were visualized using R package genoplotR (Guy et al. 2010).

### Two distinct classes of RcGTA ‘head-tail’ homologs are present in α-proteobacteria

Within 187 α-proteobacterial genomes with at least one ‘head-tail’ cluster homolog, the RcGTA-like genes are dispersed across 474 genomic regions, 245 of which contain only 1-2 RcGTA homologs (**Figure 3**). Due to significant similarity between RcGTA and genuine viral genes, some of these regions may belong to prophages unrelated to RcGTA. Indeed, 261 out of 474 regions (55%) overlap with putative prophages identified using PhiSpy (Akhter et al. 2012) (**Supplementary Figure S3**). However, PhiSpy also classified the RcGTA as a prophage, suggesting that the predictions may include other false positives. Of 889 PhiSpy-predicted prophage regions within 255 α-proteobacterial genomes, only 535 (60%) are associated with an integrase gene - a gene expected to be present in a functional prophage but not in a GTA. RcGTA-like regions are overrepresented among the predicted prophages without an integrase gene (234 out of 354 [66%] vs. 27 out of 535 [5%]), hinting that PhiSpy might not be able to distinguish GTAs from genuine prophages. Also, PhiSpy did not detect small RcGTA-like regions due to the program’s requirement of at least five consecutive genes with "viral" functional annotations in a window of 30 genes. Therefore, we examined the RcGTA-like regions, and genes within them, for additional characteristics associated with viral genes – skewed nucleotide composition (Rocha & Danchin 2002) and faster substitution rates (Drake 1999; Paterson et al. 2010) – as well as the presence of neighboring viral genes.

**Figure 3:**
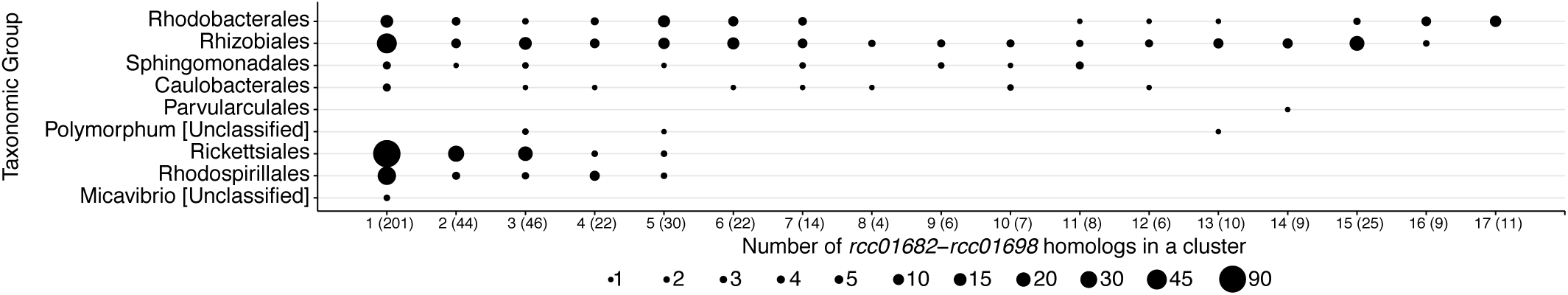
**Size variation of the RcGTA-like ‘head-tail’ clusters among α-proteobacterial genomes.** The number of homologs per genome (1-17) is shown on the X axis. The total number of genomes in each size category is shown in parentheses on the X-axis labels. The genomes are binned into taxonomic groups (arranged on Y axis). The diameter of each circle is scaled with respect to the number of genomes (see inset for scale). Only 91 of 474 clusters (19%) have at least nine RcGTA homologs. Sizes of RcGTA-like regions across several other bacterial taxonomic groups are shown in **Supplementary Figure S2**.

Genes of viral origin generally have lower fraction of Gs and Cs (GC content) than their host genomes (Rocha & Danchin 2002). However, we do not observe this trend for the RcGTA-like regions. GC content of the regions with less than nine RcGTA homologs (hereafter referred as small clusters, or SC) is, on average, equivalent to the GC content in the host (median of percent of relative change = −0.1%; pairwise Wilcoxon test; *p-*value=0.60), albeit with substantial variability across examined genomes (**Figure 4**). On the other hand, GC content of the regions with at least nine RcGTA homologs (hereafter referred as large clusters, or LC) is consistently, and in most cases significantly, higher than the GC content of their host (median of percent of relative change = 6.57%; pairwise *t* test; *p*-value < 0.001) (**Figure 4**). The elevated GC content of LCs also persists when compared to randomly sampled regions of the host genome of equivalent size (pairwise t-test; *p-value* <0.001). These observations prompted us to examine the features of LCs and SCs separately.

**Figure 4:**
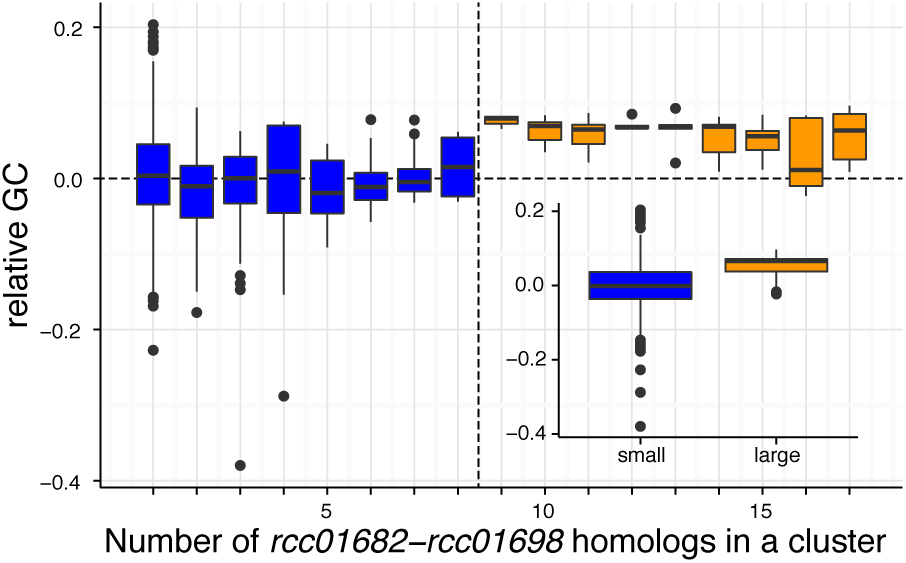
**Difference in GC content between clusters of RcGTA ‘head-tail’ gene homologs and their host chromosome.** The relative difference in GC content is calculated as (GC_GTA_-GC_host_)/GC_host_ (Y axis), and the data is presented by the number of RcGTA ‘head-tail’ homologs found in a cluster (X axis). In the box-and-whisker plots the horizontal line marks the median; the boxes extend from the 25th to 75th percentile; the whiskers encompass values at most 1.5 times away from the 25th and 75th percentile range, and any value outside of the range is shown as a dot. The vertical dashed line separates two groups of clusters: ≤ 8 genes, or small clusters (SC, in blue) and ≥9 genes, or large clusters (LC, in orange). The inset depicts the combined data from all small and large clusters.

Using the pairwise phylogenetic distances (PPD) as a proxy for substitution rates, we found that LC genes evolve significantly slower than both their SC (all of the 13 possible pairwise comparisons; Wilcoxon tests; *p-*values <0.0001) and viral homologs (14 out of 15 possible comparisons; Wilcoxon tests; *p-*values <0.005) (**Supplementary Figure S4** and **Supplementary Table S5**). While many SC genes also evolve significantly slower than their viral homologs (six out of 10 possible comparisons; Wilcoxon tests; *P-*values <0.005), the 25-75 percentile range for PPDs of SC genes often overlapped with that of the viral homologs (**Supplementary Figure S4**). Hence, SCs are more virus-like than LCs. This conjecture is supported by two additional observations. First, SCs are more frequently associated with unrelated viral genes than LCs (76 vs. 56%; **Supplementary Figure S3**). In particular, SCs are more likely to reside in a vicinity of an integrase gene than LCs (45 out of 383 SCs vs. 1 out of 91 LCs). In some cases, SCs likely belong to viral elements that are clearly not GTAs: for example, in all analyzed genomes of the *Rickettsiales'* genera *Anaplasma* and *Ehrlichia* a singleton homolog of *rcc001695* (g12) is found within 1 kb of *vir*B homologs, which in *Anaplasma phagocytophilum* belong to a virus-derived type IV secretion system (Al-Khedery et al. 2012). Second, SCs have very little conservation of flanking genes (**Supplementary Figure S5)**, while in LCs flanking genes are conserved within orders and sometimes even across larger phylogenetic distances (**Supplementary Figure S6)**.

Taken together, we hypothesize that large and small clusters represent genomic regions under different selective pressures in the host chromosome and may have separate evolutionary histories. We further hypothesize that most SCs are viral elements unrelated to RcGTA and only LCs represent RcGTA-like elements.

### RcGTA-like element was likely absent in the last common ancestor of α-proteobacteria

In what lineage did the RcGTA-like element originate? How did it propagate across α-proteobacteria? To address these questions, we examined the phylogenetic history of LCs. Within α-proteobacteria, LCs are found only within a monophyletic clade defined by node 1 in **Figure 5** (hereafter referred as Clade 1), which consists of all analyzed representatives from the orders *Rhizobiales, Caulobacterales*, *Rhodobacterales, Parvularculales* and *Sphingomonadales*. Two deeper branching α-proteobacterial clades, which include *Rickettsiales*, *Rhodospirillales,* and yet unclassified *Micavibrio* spp. and SAR 116 cluster bacteria, contain only SCs (**Figure 5**). The largest of these SCs is made up of five genes, but most are singletons (**Figures 2** and **3**). RcGTA homologs are completely absent from *Pelagibacterales* (formerly SAR11) and the basal α-proteobacterial order *Magnetococcales* (**Figure 5**), although only five genomes from these two groups were available for the analyses.

**Figure 5:**
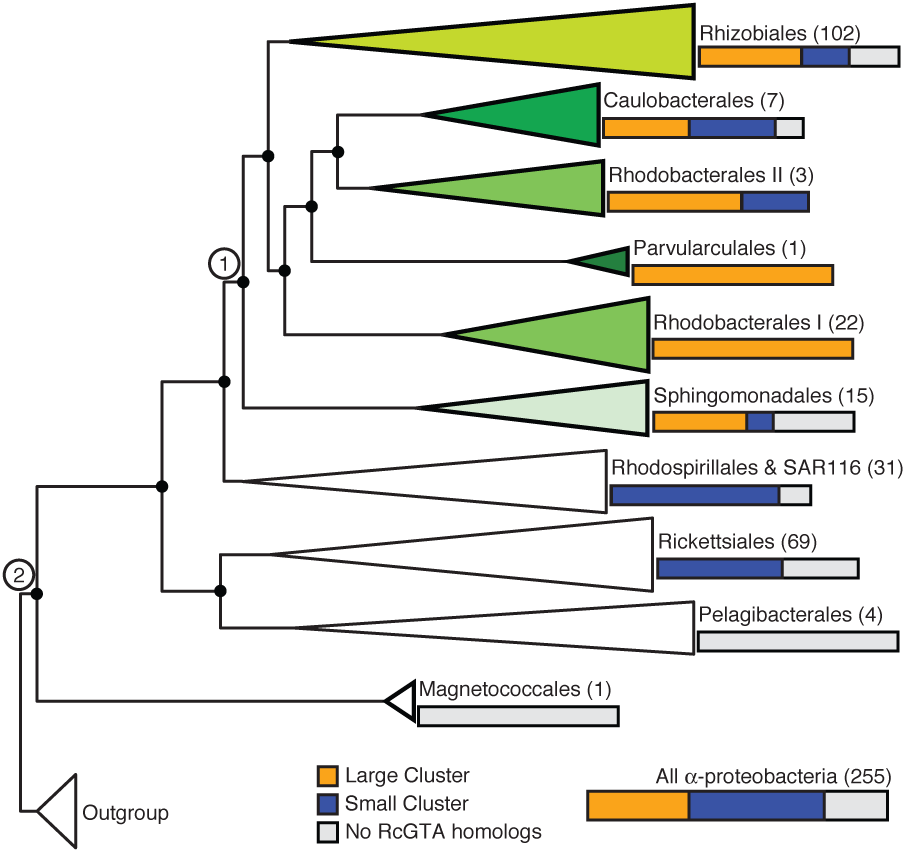
**Distribution of RcGTA homologs in α-proteobacteria.** The phylogenetic tree is a schematic version of the α-proteobacterial reference phylogeny, in which branches are collapsed at the taxonomic order level. The number of taxa in each collapsed clade is shown in parentheses. The tree is rooted using taxa from β-, γ- and δ-proteobacteria ("outgroup"). The non-collapsed version of this tree is provided in **Supplementary Figure S3**. The nodes with at least 60% bootstrap support are labeled with a black circle. Bars associated with each clade show the relative proportion of genomes that contain at least one LC (orange), only SC(s) (blue), or no detectable RcGTA homologs (gray). Nodes 1 and 2 define Clades 1 and 2 discussed in the text. Twenty-three of 150 genomes within Clade 1 and 32 out of 105 genomes outside of Clade 1 have no detectable RcGTA homologs. While relationships among the α-proteobacterial orders remain debated (Ferla et al. 2013; Gupta & Mok 2007; Lee et al. 2005; Luo 2014; Thrash et al. 2011; Williams et al. 2007), the alternative histories do not affect the inferences of this study (data not shown). Scale bar, substitutions per site.

It is possible that SCs represent decaying remnants of the previously functional GTAs. Bacterial genomes do not generally contain many pseudogenes due to genome streamlining (Lawrence et al. 2001). However, if pieces of RcGTA homologs that no longer form an ORF remain within the regions, they may still be detectable by similarity searches. Indeed, putative pseudogenes of RcGTA-like genes were detected in both large and small clusters, but only 35% of them reside in SCs (**Supplementary Figures S2, S5** and **S6**). Furthermore, only 3 of 199 SCs found in taxa outside of Clade 1 contain detectable putative pseudogenes of the RcGTA ‘head-tail’ genes (**Supplementary Figure 3**), suggesting that SCs outside of Clade 1 are probably unrelated to RcGTA.

Taking into account the observation that the fraction of genomes with no detectable RcGTA homologs is higher in taxa outside than inside of Clade 1 - 30% versus 15% of the genomes, respectively, - we hypothesize that the RcGTA-like element originate at the earliest on the branch leading to the last common ancestor of Clade 1 and not in the last common ancestor of α-proteobacteria (Clade 2 in **Figure 5**), as was previously suggested (Lang et al. 2002; Lang & Beatty 2007).

### Evolution of the RcGTA-like genome: Vertically Inherited or Horizontally Exchanged?

Did the RcGTA-like element appear on the branch leading to the last common ancestor of Clade 1, or did it spread across Clade 1 via HGT? Lang & Beatty (2007) inferred that RcGTA-like element had evolved vertically within α-proteobacteria, although only few genomes were available at the time. Within *Rhodobacterales* RcGTA-like ‘head-tail’ clusters are also predicted to be vertically inherited, but phylogenetic trees of many individual genes were poorly resolved (Hynes et al. 2016). In contrast, homologs of *rcc00171* were likely horizontally exchanged within *Rhodobacterales* (Hynes et al. 2016). To investigate the extent of HGT in the evolution of the RcGTA-like ‘head-tail’ genes across Clade 1, we reconstructed the phylogeny of LC genes and compared it to the reference tree of α-proteobacteria represented by a set of conserved α-proteobacterial genes.

Of 83 internal branches of the reference phylogeny, 42 are in disagreement with the branching order of the LC-locus phylogeny (**Figure 6**). Thirty-one of the 42 conflicts (74%) are within taxonomic orders, with 14 out of the 21 strongly supported conflicts limited to genera (**Figure 6** and **Supplementary Figure S7**). Among the nine conflicts at deep branches, eight are due to an alternative position of *Methylobacterium nodulans* ORS 2060 (**Figure 6** and **Supplementary Figure S7**), which, in addition to grouping outside of its order *Rhizobiales,* has five non-identical LCs (**Supplementary Figure S6**). Therefore, evolution of ‘head-tail’ locus is likely impacted by HGT events, but most of them have occurred between closely related taxa.

**Figure 6:**
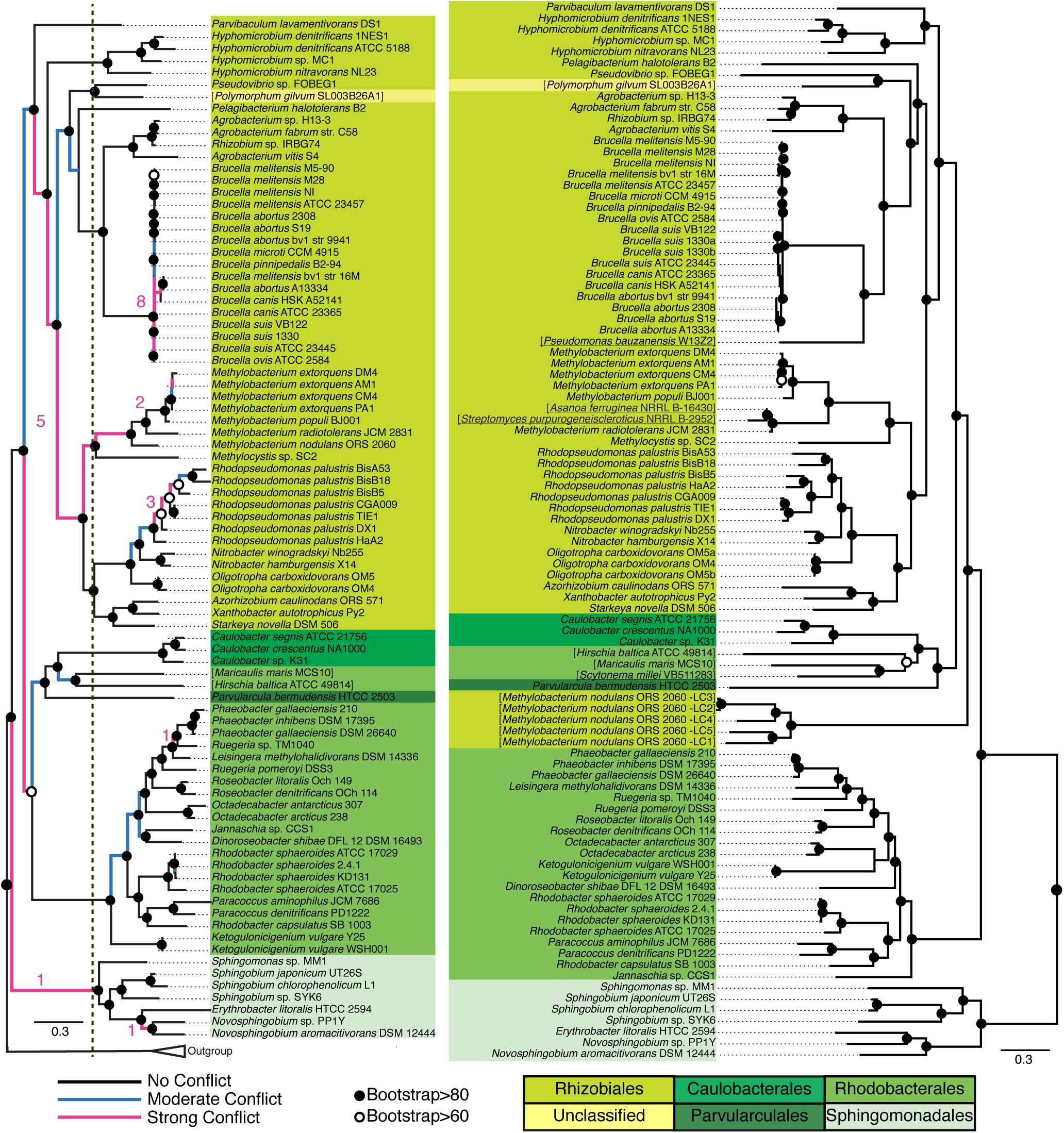
**Comparison of the phylogenetic history of large clusters (right) to the reference α-proteobacterial phylogeny (left).** The branches of the reference tree are color-coded according to the strength of conflict, as measured by an Internode <u>C</u>ertainty (IC) value (Salichos et al. 2014). IC values for all branches are shown in **Supplementary Figure S7.** The overall number of branches with strongly conflicting IC values are shown as numbers (in pink) for “deep” and “shallow” branches, as demarcated by a vertical dashed line. Fourteen of the 15 strong conflicts at shallow branches occur within genera, and five of the six strong conflicts at deep branches are due to the different position of *Methylobacterium nodulans* ORS 2060. The phylogenetic tree on the left is rooted with the same outgroup as in **Supplementary Figure S3**, while the phylogenetic tree on the right should be considered unrooted. Underlined taxa names represent four non-α-proteobacterial genomes that contain an LC (see also **Supplementary Figures S6** and **S9**). Taxa names in brackets represent lineages that branch within a taxonomic order different from the one designated by the NCBI Taxonomy database (NCBI Resource Coordinators 2017). The tree heights (*i.e.* the sum of all branch lengths of a phylogenetic tree) of the LC-locus and reference phylogenies are 31.19 and 19.39 substitutions per site, respectively, suggesting that LC locus evolves ~1.6 times faster than the conserved α-proteobacterial genes. Scale bar, substitutions per site.

Intriguingly, at least occasionally LCs have also been transferred across very large evolutionary distances. While LCs are predominantly found in α-proteobacteria, the genomes of the actinobacteria *Streptomyces purpurogeneiscleroticus* NRRL B-2952 and *Asanoa ferruginea* NRRL B-16430, the cyanobacterium *Scytonema meliloti* VB511283, and the γ-proteobacterium *Pseudomonas bauzanensis* W13Z2 also contain one LC each (**Supplementary Figures S2** and **S6**). These four non-α-proteobacterial taxa branch within the Clade 1 in three separate locations (**Figure 6**), indicating that these taxa likely acquired the LCs from the α-proteobacteria at least three times independently. Moreover, genes immediately upstream and downstream of *S. millei* and *P. bauzanensis’* LCs are homologous to the genes that flank LCs in their respective α-proteobacterial sister taxa (**Supplementary Figure S6**). These flanking genes are also of α-proteobacterial origin (**Supplementary Figure S8**), and hence were likely acquired in the same HGT events.

The duration of RcGTA-like elements’ association with bacterial genomes can also be gauged from the stability of the host chromosome regions that surround LCs. For example, highly mobile genes, such as transposable elements and some prophages, are often found in genomic islands (Juhas et al. 2009) -dynamic genomic regions that exhibit poor gene synteny even across short phylogenetic distances (Rodriguez-Valera et al. 2016). On the other hand, gene synteny of stable regions is preserved across larger evolutionary distances, although it is still subject to decay due to gene rearrangements (Brilli et al. 2013; Rocha 2006). LCs found in the same taxonomic order are generally flanked by the same, conserved α-proteobacterial genes (**Supplementary Figure S6**), hinting that they might be part of stable regions of bacterial chromosomes and not within genomic islands. We modeled gene order decay upstream and downstream of LC loci as a function of phylogenetic distance (Rocha 2006) and compared the conservation of genes flanking LCs to that of a transposable element from IS911/IS3 family and two operons of conserved housekeeping genes, the ribosomal protein operon and the ATP synthase operon. The IS911/IS3 element can undergo non-targeted integration (Chandler et al. 2015), and thus, genes adjoining these transposons are not expected to be homologous even across closely related taxa. In contrast, gene order near presumably non-mobile housekeeping operons are expected to be much more conserved across time. Indeed, we found that the regions surrounding the transposable element are too variable to even fit the model, while the decay of gene synteny near LCs is similar to that of the two operons (**Supplementary Figure S9**). Therefore, it is unlikely that RcGTA-like elements reside in dynamic regions of their host genomes.

Although sequence divergence of the remaining four loci of the RcGTA genome makes it difficult to track their evolutionary history, in 66 α-proteobacterial genomes with detected homologs of *rcc00171* and *rcc00555*, 22 LCs from Rhizobiales and Rhodobacterales have *rcc00171* homologs at their 3’ end and 6 of the 22 LCs additionally have *rcc00555* immediately downstream of an *rcc00171* homolog (**Supplementary Figure S6**). This anecdotal evidence suggests that in the past RcGTA-like elements may have carried at least some of the genes in one locus with the ‘head-tail’ genes, and that the division of the GTA genome into multiple loci may have happened after the bacterium-GTA association was already established.

Taken together, our analyses suggest a complex evolutionary history of the RcGTA-like elements. The congruence of the reference and the LC-locus phylogenies at taxonomic order and family levels and conservation of LC-flanking genes within orders suggest that RcGTA-like element was likely acquired before the contemporary orders within Clade 1 have diversified, and since that time the element co-existed with the clade members. However, the congruence does not automatically equate to vertical inheritance, since even genes highly conserved across α-proteobacteria are not immune to HGT and thus the reference phylogeny may not represent the strictly vertical history of α-proteobacteria (Gogarten et al. 2002; Andam et al. 2010). Indeed, strong phylogenetic conflicts between the reference and LC locus phylogenies of Clade 1 at both deep and shallow branches, and presence of LCs in several taxa outside of the α-proteobacteria indicate that HGT has been shaping the evolution of RcGTA-like element to a non-negligible degree. Since the majority of the strongly supported conflicts are found within genera, HGT probably played a larger role in the recent evolution of RcGTA-like elements than in their dissemination across Clade 1.

## 4 Discussion

Our survey of RcGTA homologs in the genomes of bacteria and bacterial viruses provide new insights into fascinatingly intertwined evolutionary histories of virus-like elements and their bacterial hosts. Although we infer that the RcGTA-like element is not as ancient as the class α-proteobacteria itself, RcGTA homologs are ubiquitous across a clade that spans multiple α-proteobacterial orders (Clade 1 in **Figure 5**) and, according to molecular clock estimates, has originated between 777 Mya (Luo et al. 2013) and 1710 Mya (with a 95% CI of 1977-1402 Ma; Gregory Fournier, pers. comm.) Therefore, either RcGTA-like element has been associated with a group of α-proteobacteria for hundreds of millions of years, or it is a vestige of a very successful virus capable of inter-species and inter-order infections. The latter scenario is supported by experimentally observed long-range HGT via GTAs produced by *Roseovirus nubinhibens* ISM (Mcdaniel et al. 2010). However, lack of CRISPR repeats that match RcGTA genome, absence of closely-related contemporary viruses in GenBank, and order-level congruence of the evolutionary histories of RcGTA and conserved α-proteobacterial genes point against the GTA being derived from a broad host-range virus that recently invaded this clade. Instead, our data are more compatible with a long-term association between a GTA and its bacterial host.

Over this time, the GTA genome has gone through extensive changes, as hinted by the putative HGT events within taxonomic orders, presence of likely pseudogenized RcGTA homologs within a few putative GTAs, and division of the RcGTA genome into multiple loci.

Could these modifications simply mean that this RcGTA-like element is just a defective prophage (Redfield 2001; Solioz & Marrs 1977) not yet purged from the genomes? Since bacterial genomes are generally under deletion bias of unnecessary DNA (Mira et al. 2001), conservation of “junk” regions across over millions of years is unexpected. Moreover, given the universal mutation bias towards AT-rich DNA (Hershberg & Petrov 2010), over such prolonged time we would expect the defective prophage region to become more AT-rich than the host DNA, and to accumulate mutations evenly across non-synonymous and synonymous sites of ORFs (Andersson & Andersson 2001). Yet, genomic regions that encode LCs are not AT-richer than the rest of the host chromosomes, and, at least across *Rhodobacterales*, RcGTA homologs do not exhibit increased rates of non-synonymous substitutions expected of pseudogenes (Lang et al. 2012), suggesting that LCs are unlikely to be decaying prophages.

Surprisingly, RcGTA-like gene clusters have a significantly elevated % G+C than chromosomes of their hosts. Evolution of GC content and its variation across and within the prokaryotic genomes is an unsolved puzzle, with many possibilities invoked to explain it (Hildebrand et al. 2010; Lassalle et al. 2015; Rocha & Feil 2010). In one hypothesis, AT-richness of highly expressed genes is explained by accumulation of additional mutations during the single-stranded DNA state required for transcription (Hildebrand et al. 2010). Since RcGTA-producing cells lyse and thus leave no progeny, the GTA genes are never expressed by cells that reproduce. Hence, it is tempting to speculate that selection for reduced gene expression may have played a role in driving the GC content of the ‘head-tail’ locus higher than that of the expressed host genes. Future testing of this hypothesis and exploring other possible explanations are necessary to account for the aberrant GC content of the RcGTA-like regions within Clade 1 genomes.

But if RcGTA-like elements are indeed functional, why do we observe putatively pseudogenized and apparently incomplete gene repertoires of many LCs when compared to the RcGTA genome? Perhaps, genomes of these elements are also split into different pieces, have some components replaced from sources unrelated to the RcGTA, or have evolved to a different functionality than RcGTA. For example, GTA gene expression and GTA particle release was demonstrated for *Ruegeria pomeroyi* DSS-3 (Biers et al. 2008) and *Rhodovulum sulfidophilum* (Nagao et al. 2015), suggesting that GTA is functional in these lineages. However, the *R. pomeroyi* and *R. sulfidophilum* genome contain no detectable homologs of *g1*, and this gene is essential for RcGTA production in *R. capsulatus* (Hynes et al. 2016). Therefore, even presumably “incomplete” LCs may encode functional GTAs – a conjecture that awaits experimental demonstrations.

With the evidence pointing against the *decaying* or a *selfish* nature of the RcGTA-like elements, and given that cells that produce RcGTA lyse (Westbye et al. 2013) and, therefore, the selection for the maintenance of RcGTA cannot occur at a level of an organism, earlier suggestions that GTA maintenance is due to some *population*-level benefits (Lang et al. 2012) remain most plausible explanation of persistence of RcGTA-like elements in the Clade 1 genomes. However, specific advantage(s) associated with acquisition of ~4kb pieces of random DNA via GTA remain to be elucidated. Among the proposed drivers of GTA maintenance are effective exchange of useful genes among the population of heterogeneous cells (Smillie et al. 2011), improved response to environmental stressors, such as DNA damage (Marrs et al. 1977; Brimacombe et al. 2015) and nutritional deficits (Lang et al. 2012). Possibilities unrelated to GTA-mediated DNA delivery, such as protection of the host population against infection by other viruses via the superinfection immunity mechanism (Díaz-Muñoz 2017), and increase of the host population resilience against various environmental stressors such as antibiotics (Wang et al. 2010), should also be considered. Perhaps, there is no single benefit associated with RcGTA-like elements across all Clade 1 taxa. Instead, different lineages may experience selection pressures unique to their ecological niches.

Interestingly, GTA-associated benefits have been proposed to drive evolutionary innovations of cellular lifeforms that go beyond population level. For example, it has been hypothesized that abundance and mixed ancestry of α-proteobacterial genes in eukaryotic genomes is due to the delivery of such genes by RcGTA-like elements into proto-eukaryote genome, and that such “seeding” facilitated the later integration of mitochondrial progenitor and the proto-eukaryotic host (Richards & Archibald 2011). Our study, however, does not support this hypothesis. First, based on the estimated age of the Clade 1 (777-1710 Mya) and the appearance of eukaryotic, and hence mitochondria-containing, organisms in the fossil record (1600 and possibly 1800 Mya; Knoll 2014), the RcGTA-like system likely originated after the eukaryogenesis took place. Second, α-proteobacterial lineage(s) suggested to have given rise to mitochondrial progenitor (Brindefalk et al. 2011; Thrash et al. 2011; Wang & Wu 2015) appear to lack LCs and therefore are unlikely to have had RcGTA-like elements. In another intriguing hypothesis, an unrelated GTA in *Bartonella* has been proposed to have facilitated adaptive radiation within the genus (Guy et al. 2013). Perhaps, acquisition of the RcGTA-like element by the last common ancestor of Clade 1 taxa represent a similar innovation that led to diversification of this clade and success of its members in a variety of ecological settings that span soil, freshwater, marine, waste water, wetland, and eukaryotic intracellular and extracellular environments.

Despite the appearance that bacteria maintain GTA genes for the potential contribution to their own evolutionary success, there is an undeniable shared evolutionary history between RcGTA-like elements and *bona fide* viruses. Presence of homologs for most RcGTA-like genes in viral genomes, as well as similarity in the organization of the ‘head-tail’ cluster and corresponding region of a typical siphovirus genome (Huang et al. 2011), led to a prevailing hypothesis that RcGTA originated in an initial co-option of a virus and evolved in subsequent modification of the progenitor prophage genome via vertical descent and HGT (Lang et al. 2017). Our data are compatible with this scenario, and further suggest an even larger role of HGT in the evolution of RcGTA-like elements than currently acknowledged, shaping GTAs both within α-proteobacterial orders, but also at least occasionally transferring RcGTA-like regions across the higher-order α-proteobacterial taxa and even to different phyla. Yet, there are hints of even more entangled connection between RcGTA-like genes found in cellular and viral genomes: a few viruses that infect *Paracoccus*, *Roseobacter*, *Caulobacter*, and *Rhodobacter* spp., and a yet unclassified marine *Rhizobiales* str. JL001 bacterium, carry RcGTA-like genes that are phylogenetically placed within their cellular homologs (Zhan et al. 2016), suggesting that some viruses acquired genes from the cellular RcGTA-like regions. Given recent propositions of (*i*) cellular origin of even such “hallmark” viral genes like those encoding capsid proteins (Krupovic & Koonin 2017), (*ii*) the possibly virus-independent origin of cellular microcompartments that resemble viral structures (Bobik et al. 2015), and (*iii*) the potentially primordial origin of type VI secretion system that resembles a bacteriophage tail (Böck et al. 2017), we should not exclude a possibility that genomic regions encoding RcGTA-like nanostructures did not originate by a co-option of a prophage and its subsequent modification, but instead are a mosaic originally assembled within bacterial genomes from the available cellular and viral parts.

## Acknowledgements

We would like to thank Andrew S. Lang, J. Thomas Beatty and Rosemary J. Redfield for numerous stimulating discussions regarding GTA evolution.

## Author Contributions

MS and OZ designed the study. MS and SMS performed the analyses. MS, SMS, and OZ wrote the manuscript. All authors have seen and approved the final manuscript.

## Data Availability

Amino acid sequences of RcGTA homologs in bacterial and viral genomes; amino acid sequence alignments of RcGTA homologs from the LCs, SCs, and viruses; concatenated alignment of 99 conserved α-proteobacterial genes and RcGTA homologs from LCs; and discussed phylogenetic trees in Newick format are available via FigShare at DOI 10.6084/m9.figshare.5406733.

## Supplementary Data

**SupplementaryFigures.pdf**: Supplementary Figures 1-9

**SupplementaryTables.pdf:** Supplementary Tables 1-5

